# Loss of the GARP but not EARP protein complex drives Golgi sterol overload during dendrite remodeling

**DOI:** 10.1101/2021.09.29.462255

**Authors:** Caitlin O’Brien, Susan H. Younger, Lily Yeh Jan, Yuh Nung Jan

## Abstract

Membrane trafficking is essential for sculpting neuronal morphology. The GARP and EARP complexes are conserved tethers that regulate vesicle trafficking in the secretory and endolysosomal pathways, respectively. Both complexes contain the Vps51, Vps52, and Vps53 proteins, and a complex-specific protein: Vps54 in GARP and Vps50 in EARP. In *Drosophila*, we find that both complexes are required for dendrite morphogenesis during developmental remodeling of multidendritic class IV da (c4da) neurons. Having found that sterol accumulates at the trans-Golgi network (TGN) in *Vps54*^*KO/KO*^ neurons, we investigated genes that regulate sterols and related lipids at the TGN. Overexpression of oxysterol binding protein (Osbp) or knockdown of the PI4K *four wheel drive (fwd)* exacerbates the *Vps54*^*KO/KO*^ phenotype, whereas eliminating one allele of *Osbp* rescues it, suggesting that excess sterol accumulation at the TGN is, in part, responsible for inhibiting dendrite regrowth. These findings distinguish the GARP and EARP complexes in neurodevelopment and implicate vesicle trafficking and lipid transfer pathways in dendrite morphogenesis.

## Introduction

Proper wiring of the nervous system depends on the development and maintenance of complex polarized neuronal morphologies. Neurons initially establish their architectures during embryogenesis, which is followed by an extended post-embryonic period during which excess branches are pruned and remodeled into their mature states (Stiles and Jernigan, 2010). Membrane trafficking pathways are essential to establishing and sculpting neuronal morphology during development (Winkle and Gupton, 2016). Secretory vesicles are essential sources of membrane for neurite outgrowth (Vega and Hsu, 2001) and dendrite development is particularly sensitive to inhibition of secretory trafficking (Ye *et al*., 2007). Endocytosis and recycling pathways are also required for growth factor-mediated branching of dendrites (Lazo *et al*., 2013) and axons (Ascano *et al*., 2009). Both the secretory (Wang *et al*., 2017; Wang, *et al*., 2018) and endocytic (Kanamori *et al*., 2015; Krämer, *et al*. 2019) pathways also play important roles during developmental pruning and remodeling of dendrites. Numerous mutations in regulators of membrane trafficking are associated with neurodevelopmental disorders (Ouyang *et al*., 2013; Ivanova *et al*., 2017; Marin-Valencia *et al*., 2017; Passemard *et al*., 2017), highlighting the importance of these pathways in proper nervous system development.

The closely related GARP (*G*olgi-*A*ssociated *R*etrograde *P*rotein) and EARP (*E*ndosome-*A*ssociated *R*ecycling *P*rotein) complexes are conserved membrane tethers that function in the secretory and endolysosomal pathways. They share the common subunits Vps51, Vps52 and Vps53 (Vps for *v*acuolar *p*rotein *s*orting) (Fig. 1a). This core interacts with either Vps54 to form the GARP complex (Conibear and Stevens, 2000; Siniossoglou and Pelham, 2002; Reggiori *et al*., 2003) or Vps50 to form the EARP complex (Schindler *et al*., 2015). The GARP complex primarily localizes to the trans-Golgi network (TGN) where it tethers endosomes and facilitates SNARE complex formation for the retrograde delivery of cargo to the Golgi (Perez-Victoria and Bonifacino, 2009). The GARP complex is required for proper sorting of various cargos, including the lysosomal hydrolases (Pérez-Victoria *et al*. 2008), and for secretion of GPI-linked proteins (Hirata *et al*., 2015). The more recently described EARP complex associates with early endosomes and facilitates Rab4-dependent cargo recycling (Gillingham *et al*., 2014; Schindler *et al*., 2015), as well as Rab2-dependent sorting into dense-core vesicles (Topalidou *et al*., 2016).

**Figure 1.**
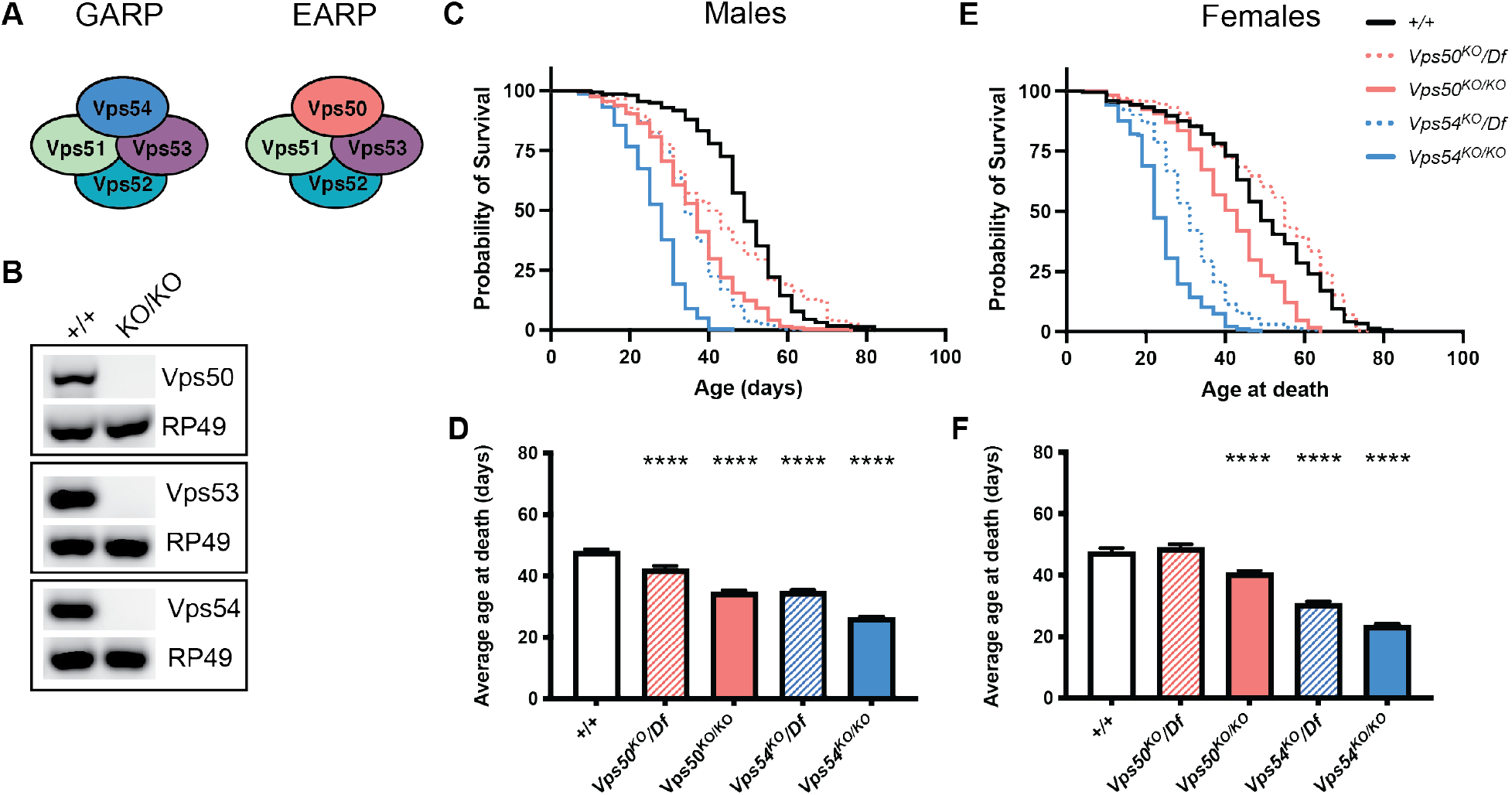
Reduced lifespan of GARP knockout flies. A) Cartoon depicting the GARP and EARP complexes. B) RT-PCR from control (+/+) and KOs. RNA was extracted from whole larvae, reverse transcribed, and equal amounts of cDNA were used for RT-PCR. RP49 is used as a reference gene. C) Survival curves and D) average age at death ± SEM for male flies of the indicated genotypes. N > 200/genotype. Survival curves were analyzed by Log-Rank Mantel-Cox test with Bonferroni multiple comparisons correction. ****p < 0.0001 for all genotypes compared to control except *Vps50*^*KO*^*/Df* (not significant – n.s.). Average age at death analyzed by one-way ANOVA. **** p < 0.0001. E) Survival curves and F) average age at death ± SEM for female flies of the indicated genotypes. N > 188/genotype. Survival curves were analyzed by Log-Rank Mantel-Cox test with Bonferroni multiple comparisons correction. ****p < 0.0001 for all genotypes compared to control except *Vps50*^*KO*^*/Df* (n.s.). N > 188/genotype.

Several neurodevelopmental disorders are associated with mutations in GARP and EARP subunits. Mutations in the core components *Vps51* (Gershlick *et al*., 2018) and *Vps53* (Feinstein *et al*., 2014; Hady-Cohen *et al*., 2018) have been identified in patients who suffer from profound developmental delays and progressive postnatal microcephaly. Mutations in *Vps50* have been linked to neural tube defects (Shi *et al*., 2019). These studies underscore the importance of the GARP and EARP complexes in neurons, prompting our study to examine their function during neuronal development. The dendritic arborization (da) sensory neurons in *Drosophila melanogaster* are a well characterized model to study dendrite morphogenesis (Grueber, *et al*., 2002). The c4da neurons establish complex larval dendritic arbors, which then undergo developmental pruning and regrowth to their mature adult forms during pupation (Kuo, *et al*., 2005; Williams and Truman, 2005; Shimono *et al*., 2009), making them amenable to study various aspects of neurodevelopment.

Cholesterol is an important component of cellular membranes, regulating membrane fluidity and protein sorting (Ikonen, 2008; Lippincott-Schwartz and Phair, 2010). While most cells can either synthesize endogenous sterols or obtain them from dietary sources, *Drosophila melanogaster* is a sterol auxotroph – they lack the ability to synthesize sterols – and must obtain sterols entirely from dietary sources (Clayton, 1964). *Drosophila* is therefore an excellent model to study the uptake and transport of sterol between organelles. Dietary sterols packaged with low-density lipoproteins (LDL) are endocytosed after binding the LDL receptor. The Niemann Pick proteins NPC1 and 2 then coordinate the non-vesicular transfer of sterols from the endolysosomal lumen to the ER through interorganelle contact sites (Infante *et al*., 2008; Höglinger *et al*., 2019). Both yeast and mammalian cells lacking the core GARP/EARP components accumulate sterol in lysosomes due to missorting of NPC2 (Fröhlich *et al*., 2015; Wei *et al*., 2017). Once in the secretory pathway, sterols are tightly regulated as the flow of cargo through the Golgi is particularly sensitive to sterol levels. Sterol depletion inhibits secretory vesicle budding from the TGN (Wang, Thiele and Huttner, 2000), while sterol overload strongly inhibits transport of the model secretory cargo VSV-G (Stüven *et al*., 2003). During neuronal development, either depletion or overload of sterols can decrease dendrite and axonal branching (Fan *et al*., 2002; Ko *et al*., 2005).

In this study, we generated CRISPR knockout flies for shared and complex-specific genes of the GARP and EARP complexes and demonstrate a role for both complexes in the development of adult c4da neuron arbors. Sterol accumulates in neurons lacking the GARP (*Vps54*^*KO/KO*^), but not EARP (*Vps50*^*KO/KO*^), complex during regrowth after developmental pruning. Unexpectedly, we find sterol accumulating at the TGN rather than lysosomes in GARP-deficient neurons. Altering the transport or availability of sterol and related lipids at the TGN modulates GARP null phenotypes. In particular, overexpressing *oxysterol binding protein (Osbp)* or knocking down the PI4P kinase, *four wheel drive (fwd)*, exacerbates the dendrite regrowth phenotype in *Vps54*^*KO/KO*^ neurons, while haploinsufficiency of *Osbp* rescues it.

## Results

### Reduced lifespan of GARP knockout flies

In mice, homozygous null mutants of *Vps52* and *Vps54* are lethal at early embryonic stages (Schmitt-John *et al*., 2005; Sugimoto *et al*., 2012), limiting the study of the GARP and EARP complexes in these models. To overcome these challenges, we made use of the genetic toolbox available in *Drosophila*. To study the role of the GARP and EARP complexes, we generated knockouts (KO) of the EARP-specific *Vps50*, the shared core component *Vps53*, and the GARP-specific *Vps54* (also known as *scattered* in flies), by using CRISPR/Cas9 gene editing to replace the entire coding sequences of each gene (see Fig. S1 for schematic and genotyping). We confirmed by RT-PCR that expression of each gene targeted for KO was eliminated (Fig. 1 b). While we could not find antibodies to determine Vps50 or Vps53 protein levels in *Drosophila*, we were able to confirm that Vps54 protein was absent from *Vps54*^*KO/KO*^ larvae (Fig. S1 h).

In flies, homozygous knockouts of *Vps53* (*Vps53*^*KO/KO*^) survived the larval stages but died during pupation. Ubiquitous expression of UAS-Vps53 using either *tubulin*-Gal4 or *daughterless*-Gal4 allowed for survival of *Vps53*^*KO/KO*^ flies to adulthood. In contrast to mice, homozygous knockout flies of the complex-specific components, *Vps50*^*KO/KO*^ or *Vps54*^*KO/KO*^, were viable to adulthood. Loss of the GARP-specific Vps54 (*Vps54*^*KO/KO*^) reduced lifespan in both males and females (Fig. 1, c–f). Control males lived an average of 46.7 ± 0.9 days and a maximum of 82 days, whereas *Vps54*^*KO/KO*^ male flies lived on average only 24.6 ± 0.6 days and a maximum of 46 days. Control female flies lived an average of 47.6 ± 1.2 days and a maximum age of 82 days, whereas *Vps54*^*KO/KO*^ females lived an average of only 23.7 ± 0.6 days and a maximum age of 49 days. We confirmed these results by crossing *Vps54*^*KO*^ flies to a chromosomal deficiency (*Df(2L)Exel8022*). Loss of the EARP-specific component *Vps50* did not consistently reduce lifespan across genotypes.

The GARP complex has also been implicated in spermiogenesis in both mice and flies (Castrillon *et al*., 1993; Schmitt-John *et al*., 2005). In fact, the name for the *Drosophila* homolog of Vps54, *scattered*, refers to the disorganized, scattered organization of nuclei in developing spermatids. To further characterize these knockouts, we therefore also tested fertility of male files. *Vps54*^*KO/KO*^ and *Vps54*^*K*^*/Df* males, like the *scat*^*1/1*^ null males were sterile. In contrast, *Vps50*^*KO/KO*^ and *Vps50*^*KO*^*/Df* males were fertile.

### Loss of either the EARP or GARP complex impairs arborization of adult neurons

To determine how knockout of the EARP and GARP complexes may affect neuron development, we first examined the overall morphology of c4da neurons in adult pharate flies. To circumvent difficulties owing to the adult lethality of *Vps53*^*KO/KO*^, we used MARCM (<u>m</u>osaic <u>a</u>nalysis with a <u>r</u>epressible <u>c</u>ell <u>m</u>arker) (Lee and Luo, 1999) to generate homozygous knockout clones in the viable heterozygous flies, to evaluate the role of *Vps53* in neuronal morphology. Dendrite arbors in *Vps53*^*KO/KO*^ clones were only about a third of the total length of controls (Fig. 2, a and b), revealing a cell autonomous requirement of *Vps53* for dendritic morphology. The arbors of *Vps53*^*KO/KO*^ clones were also less complex and contained fewer total branches (Fig. 2, c and d). The cell-autonomous involvement of *Vps53* was validated by showing that the dendrite branch length and number in the *Vps53*^*KO/KO*^ clones can be rescued by using the class IV specific *ppk*-Gal4 to drive expression of wildtype Vps53 protein.

**Figure 2.**
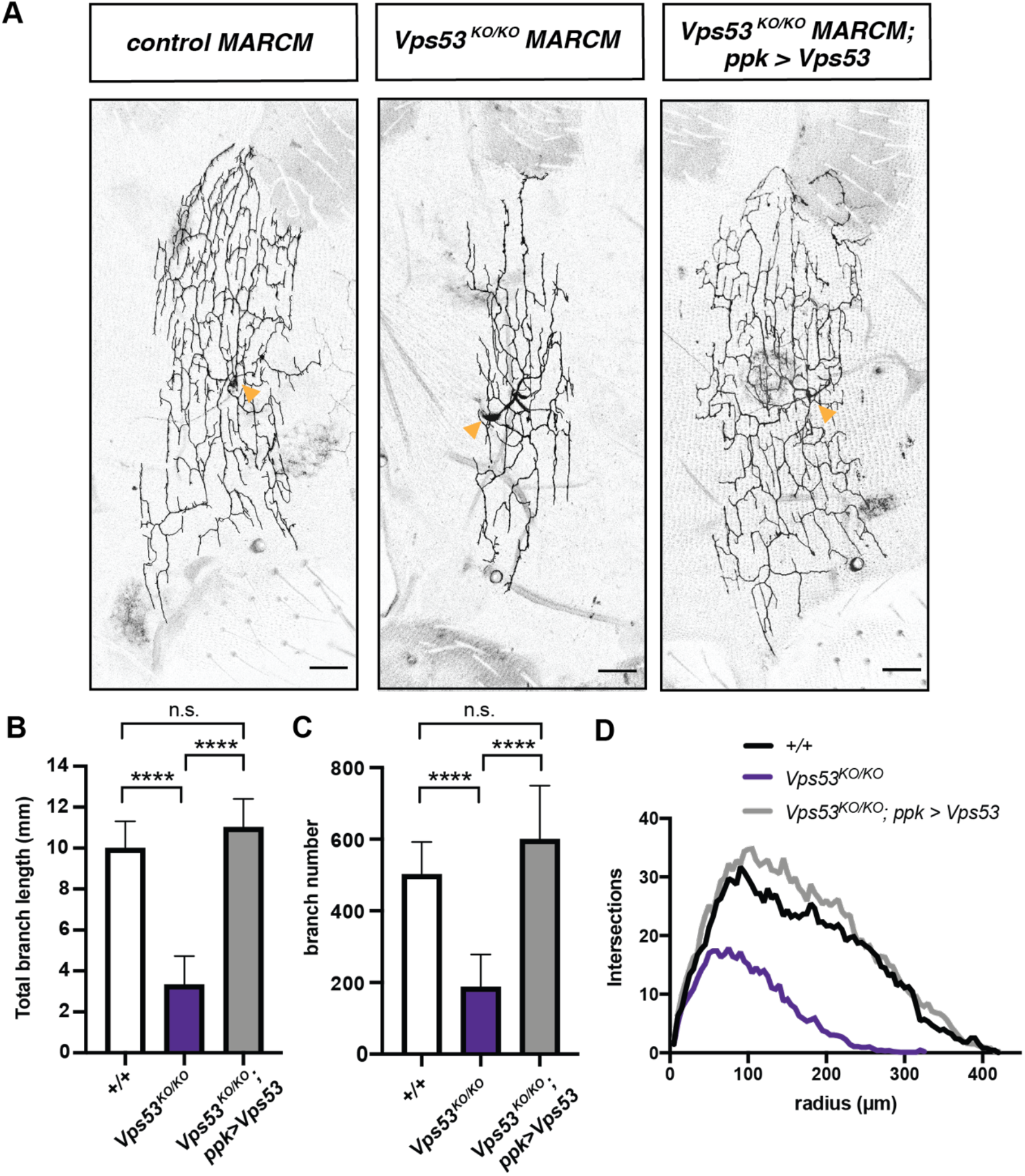
Neuron-specific knockout of *Vps53* results in smaller dendritic arbors. A) Representative maximum z-projections of MARCM control FRT^40A^, *Vps53*^*KO/KO*^, and *Vps53*^*KO/KO*^; ppk>Vps53 class IV da neuron clones. Images were collected from 7 day old male pharate adults. Yellow arrows point to the soma. Scale bar = 50μm. B) Quantification of total dendrite branch length and C) total branch number presented as mean ± standard deviation. Both total dendrite branch length and number were analyzed by one-way ANOVA with Tukey’s post-test. **** p <0.0001. D) Sholl analysis. MARCM control FRT^40A^ n = 8; *Vps53*^*KO/KO*^ n = 13; and *Vps53*^*KO/KO*^; *ppk>Vps53* n = 11.

Given that loss of *Vps53* disrupts both the EARP and GARP complexes, we next analyzed neuronal morphology in knockouts targeting the complex-specific components, *Vps50* and *Vps54*, respectively (Fig. 3, a–d). Whole body knockout of either *Vps50* or *Vps54* reduced the total dendritic length and branch number as compared to controls. For both parameters, the GARP-specific *Vps54*^*KO*^ had a stronger effect than *Vps50*^*KO*^. These dendrite morphology defects can be rescued by expression of the respective wildtype protein in neurons via the *ppk*-Gal4, suggesting that the GARP and EARP complexes function cell autonomously to regulate dendrite morphogenesis. RNAi knockdown of EARP and GARP complex components in c4da neurons further confirmed the cell-autonomous function of these complexes on adult dendrite arborization (Fig. S2 a).

**Figure 3:**
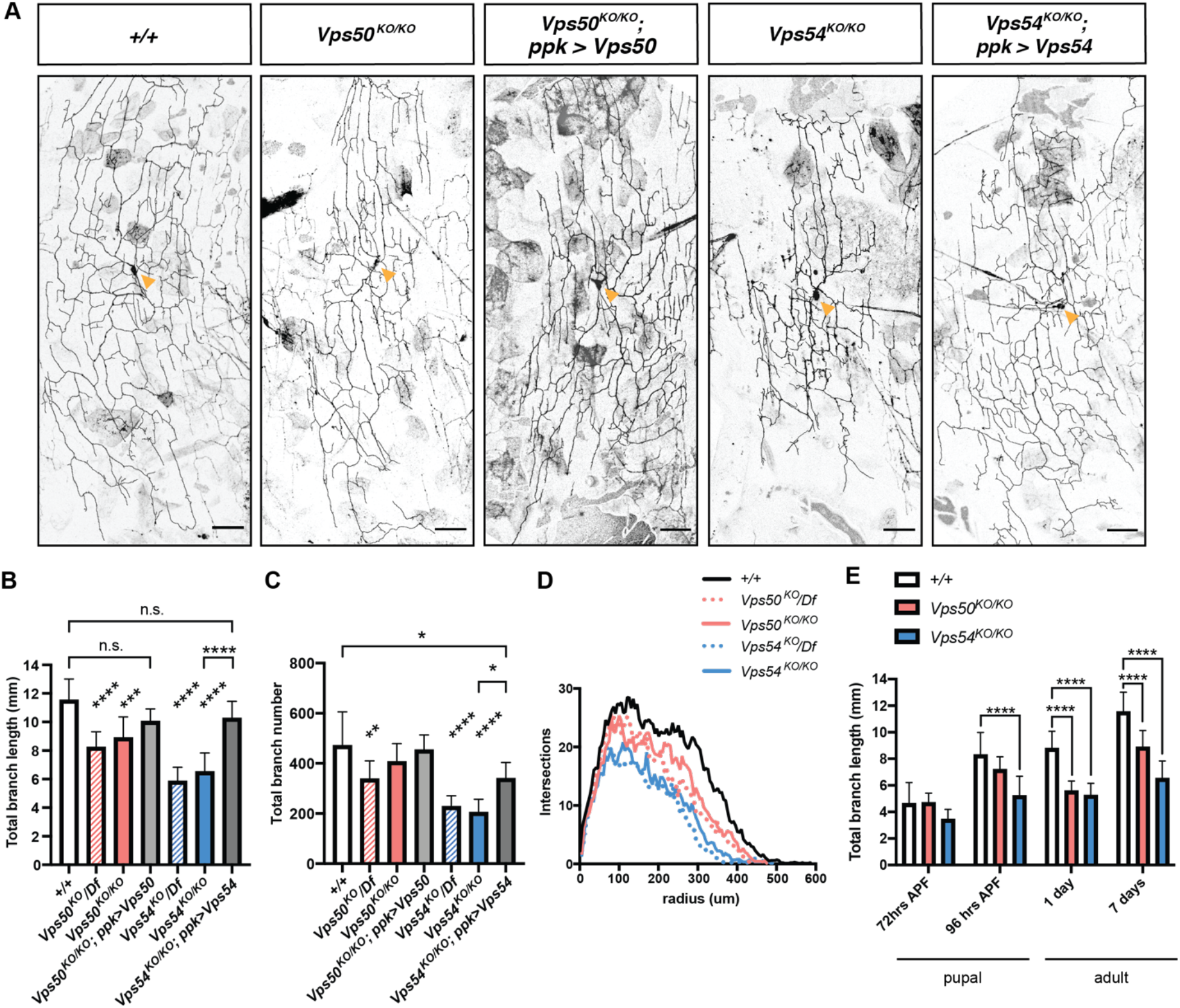
Both the GARP and EARP complexes are necessary for dendrite arborization. A) Representative maximum z-projections of class IV da neurons from 7 day old male pharate adults. Yellow arrows point to the soma. Scale bar = 50 μm. B) Quantification of total dendrite length and C) total dendrite branch number. Both total dendrite branch length and number were analyzed by one-way ANOVA with Tukey’s post-test. * p < 0.05, ** p<0.01 **** p <0.0001. D) Sholl analysis. For B-D, n = 7-12/genotype. E) Quantification of total dendrite branch length over development, +/+ n = 10/timepoint; *Vps50*^*KO/KO*^ n = 9-11/timepoint; *Vps54*^*KO/KO*^ n = 10-12/timepoint. Analyzed by two-way ANOVA with Tukey’s post-test. * p < 0.05, ** p<0.01 **** p <0.0001.

C4da neurons establish dendritic arbors initially in the larval stage. Early in pupation, the dendrites are extensively pruned and these neurons subsequently regrow a remodeled adult arbor (Kuo, *et al*., 2005; Williams and Truman, 2005; Shimono *et al*., 2009). To gain a better understanding of when the EARP and GARP complexes are required during this dynamic period of development, we analyzed dendrite morphology in both larvae and pupae. C4da neurons in *Vps50*^*KO/KO*^, *Vps53*^*KO/KO*^ or *Vps54*^*KO/KO*^ larvae were relatively spared (Fig. S2 b). There does not appear to be any significant effect of maternally contributed Vsp50 or Vps54 to dendrite growth, as neurons in larvae from homozygous knock-out mothers grew arbors comparable in size to controls. However, while regrowth of the adult arbor during pupation began normally in knockout flies, it ultimately stalled (Fig. 3 e). The *Vps54*^*KO/KO*^ neurons stopped growing just before eclosion by 96 hrs after puparium formation (APF), while the *Vps50*^*KO/KO*^ phenotype emerged slightly later in 1 day old adults.

In order to determine if the knockout phenotype was limited to dendrites, we also examined overall axon morphology. The c4da neurons project to the ventral nerve cord (VNC), where they form synapses with second order sensory neurons (Tsubouchi *et al*., 2017). We did not observe any gross changes in axon morphology in newly-eclosed 1 day old *Vps50*^*KO/KO*^ or *Vps54*^*KO/KO*^ adults, suggesting that the dendritic phenotype is related to regrowth of the arbor rather than axon retraction or growth factor withdrawal.

### Complex specific defects in secretory and endosomal organelles

Given the role of the EARP and GARP complexes in regulating specific steps in membrane trafficking, we examined various markers of the endolysosomal and secretory pathways in 1 day old adults, when knockouts of both complexes exhibited dendrite phenotypes. The number of Rab5+ early endosomes was increased in the soma of *Vps50*^*KO/KO*^ but not *Vps54*^*KO/KO*^ neurons (Fig. 4, a and b), consistent with the fact that the EARP complex facilitates cargo sorting from early endosomes to Rab4+ recycling endosomes (Schindler *et al*., 2015). In proximal dendrites, we did not observe a similar increase in the number of early endosomes in *Vps50*^*KO/KO*^ neurons (Fig. S3 a). The number of Rab7+ late endosomes was increased in the soma of *Vps54*^*KO/KO*^ but not *Vps50*^*KO/KO*^ neurons (Fig. 4, c and d), indicative of complex-specific defects in endosome populations. Dendrites are devoid of degradative lysosomes, and therefore endosomal cargo destined for degradation must be trafficked to the soma (Yap *et al*., 2018). We did not observe any significant change in Rab7+ positive endosomes in the dendrites of *Vps54*^*KO/KO*^ neurons (Fig. S3 b), suggesting their trafficking was not affected. These changes in endosomal populations did not occur in larval neurons (Fig. S3, c and d), supporting the notion that these complexes are dispensable for larval neurodevelopment.

**Figure 4.**
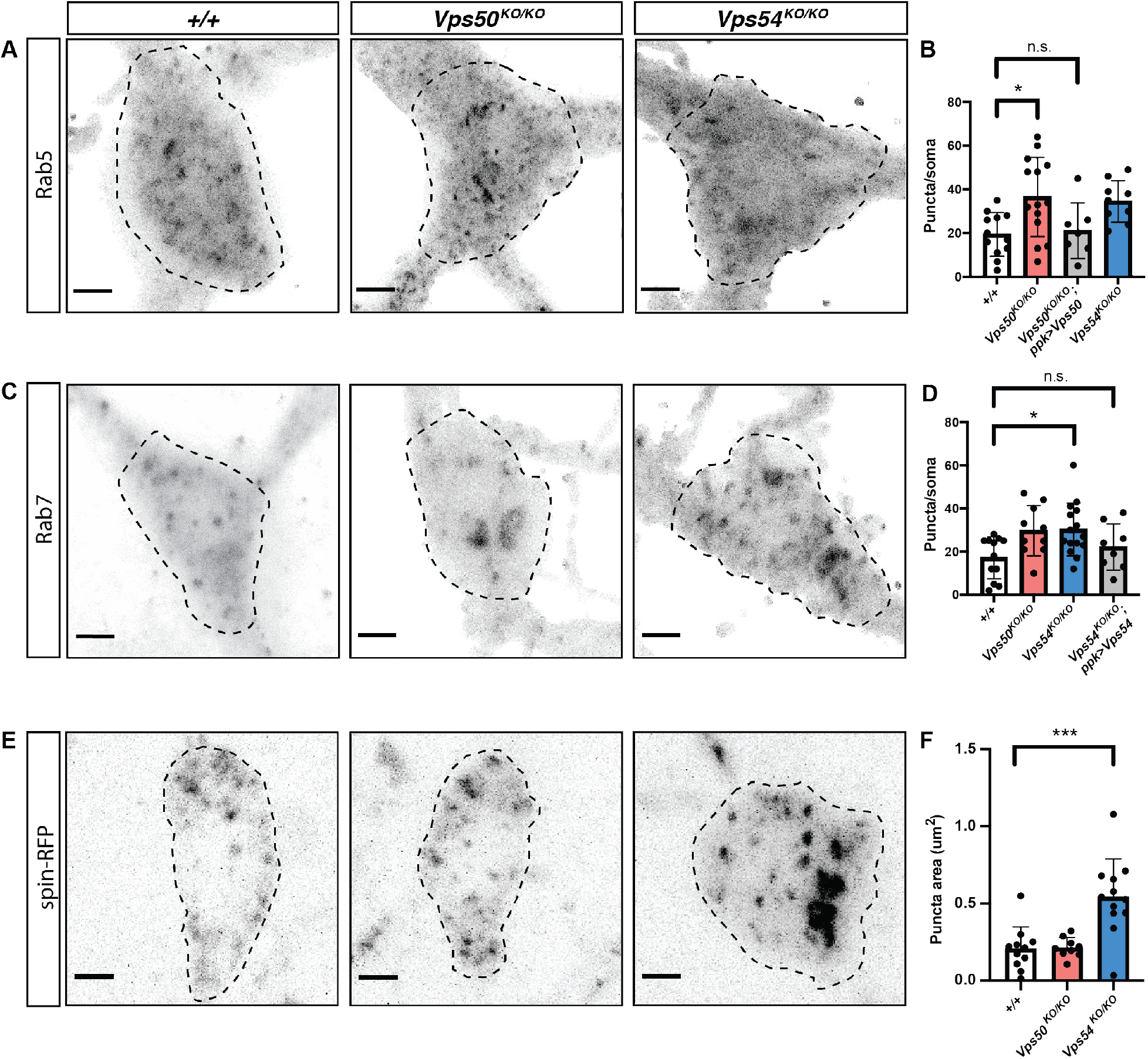
Complex-specific impairments in endosome populations. A) Maximum intensity projections of endogenous Rab5 staining in neurons from 1 day old flies. Dashed lines indicate soma area. B) Quantification of the number of Rab5 puncta/soma, n = 7-12/genotype. C) Maximum intensity projections of endogenous Rab7 staining. D) Quantification of the number of Rab7 puncta/soma, n = 8-16/genotype. E) Maximum intensity projections of spinster-RFP. F) Quantification of spin-RFP puncta area, n = 9-12/genotype. Scale bar = 2.5μm. Puncta number were analyzed by one-way ANOVA with Tukey’s post-test. * p<0.05, *** p<0.001.

We also examined the lysosomal marker *spin-RFP* and observed an expansion of this compartment in the soma of *Vps54*^*KO/KO*^, but not *Vps50*^*KO/KO*^, neurons. Lysosomal expansion can be a result of impaired cargo degradation by the resident acid hydrolases such as cathepsins. Immature forms of acid hydrolases are trafficked from the secretory pathway to lysosomes in a GARP-dependent manner (Pérez-Victoria *et al*. 2008). Upon reaching the acidic environment of the lysosome, hydrolases are processed into their mature, active forms. Therefore, we also examined the processing of cathepsin L (catL) by western blot in head lysates. We did not observe any difference in catL processing in young adult flies in *Vps54*^*KO/KO*^ or *Vps50*^*KO/KO*^ neurons (Fig. S3, e–g), suggesting that catL is successfully trafficked to acidic lysosomes in both knockout lines. Further, these results suggest that the inability of neurons to regrow their adult arbors during pupation may be independent of their lysosomal degradative capacity.

Because the GARP complex regulates retrograde traffic to the TGN, we also examined this compartment by staining for Golgin245. The number of Golgin245 puncta was increased in both the soma and proximal dendrites of *Vps54*^*KO/KO*^ but not *Vps50*^*KO/KO*^ neurons (Fig. 5, a, b, and d). If the increase in puncta number were due to fragmentation of the Golgi, we would expect the puncta to be smaller in size. However, we did not observe a significant difference in the size of Golgin245 puncta (Fig. 5, c and e), suggesting the increase in puncta number is not a result of Golgi fragmentation.

**Figure 5.**
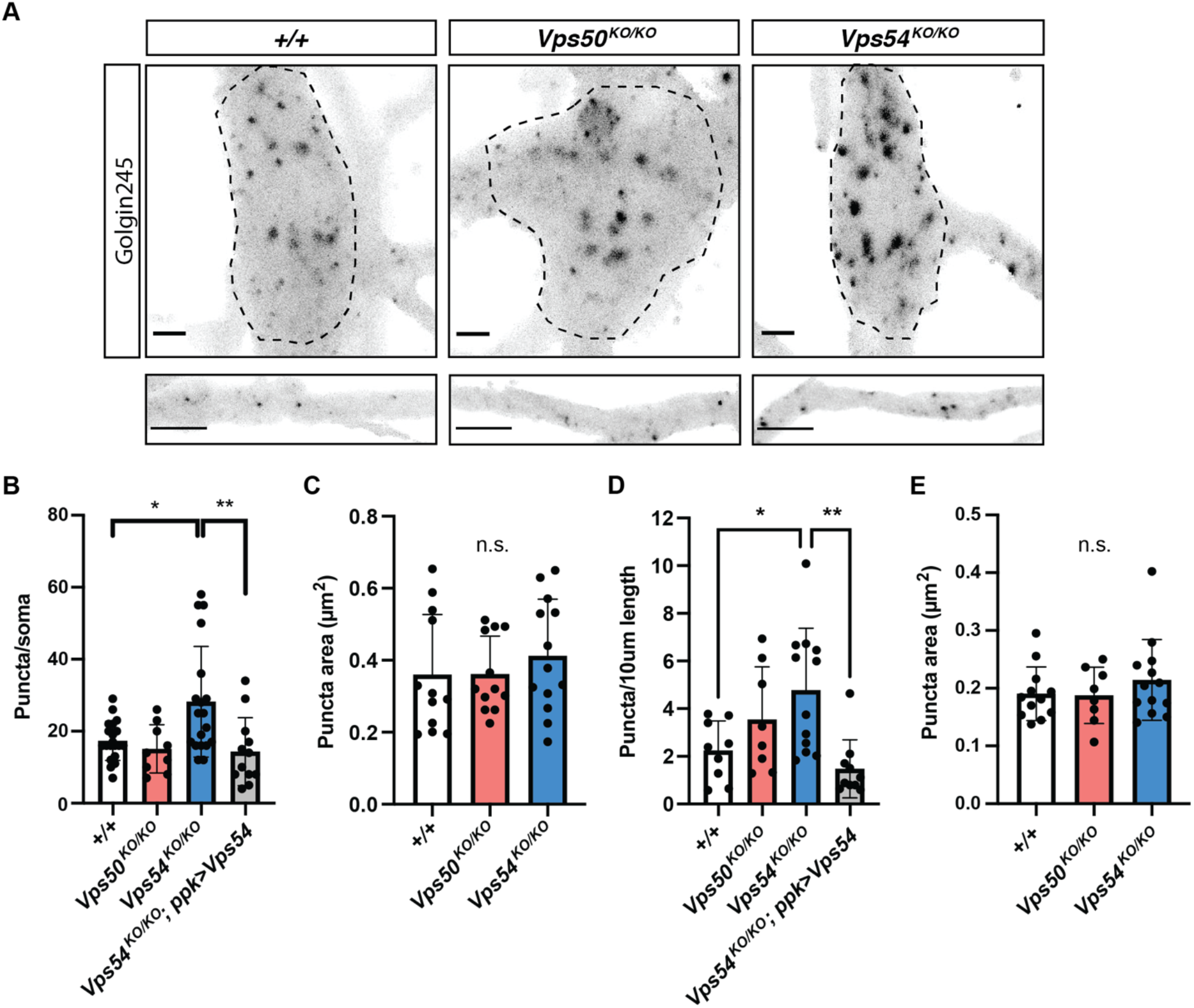
Expansion of the TGN in GARP-deficient neurons. A) Maximum intensity projections of endogenous Golgin245 staining in neurons from 1-day old flies. Top: soma, bottom: proximal dendrites. Dashed lines indicate soma area. Quantification of B) puncta/soma and C) average puncta area in soma. Quantification of D) puncta/10μm of dendrite length and E) average puncta area in dendrites. Scale bar for soma = 2.5μm. Scale bar for dendrites = 5μm. Puncta number were analyzed by one-way ANOVA with Tukey’s post-test. * p<0.05, ** p<0.01.

### Sterols accumulate at the TGN rather than endolysosomes in GARP KO neurons

Previous studies have reported accumulation of sterols in cells lacking the GARP complex (Fröhlich *et al*., 2015; Wei *et al*., 2017). We therefore sought to examine sterol levels and localization in knockout neurons. Filipin is a widely used fluorescent stain that binds to free sterols. *Vps54*^*KO/KO*^ neurons exhibited strong internal filipin staining compared to controls, which was rescued by expression of wildtype Vps54 (Fig. 6, a and b). Sterol accumulation in GARP deficient neurons appeared to be transient and to correlate with the emergence of the dendrite morphology defect in *Vps54*^*KO/KO*^ neurons (Fig. 6 c). Filipin staining was comparable between *Vps54*^*KO/KO*^ and control neurons in 3rd instar larvae but increased in the *Vps54*^*KO/KO*^ neurons during pupation, peaking at 96 hrs APF. *Vps50*^*KO/KO*^ neurons, however, showed filipin staining comparable to controls (Fig. 6 b and Fig. S4 a), indicating that the GARP, but not EARP complex, plays a role in sterol processing.

**Figure 6.**
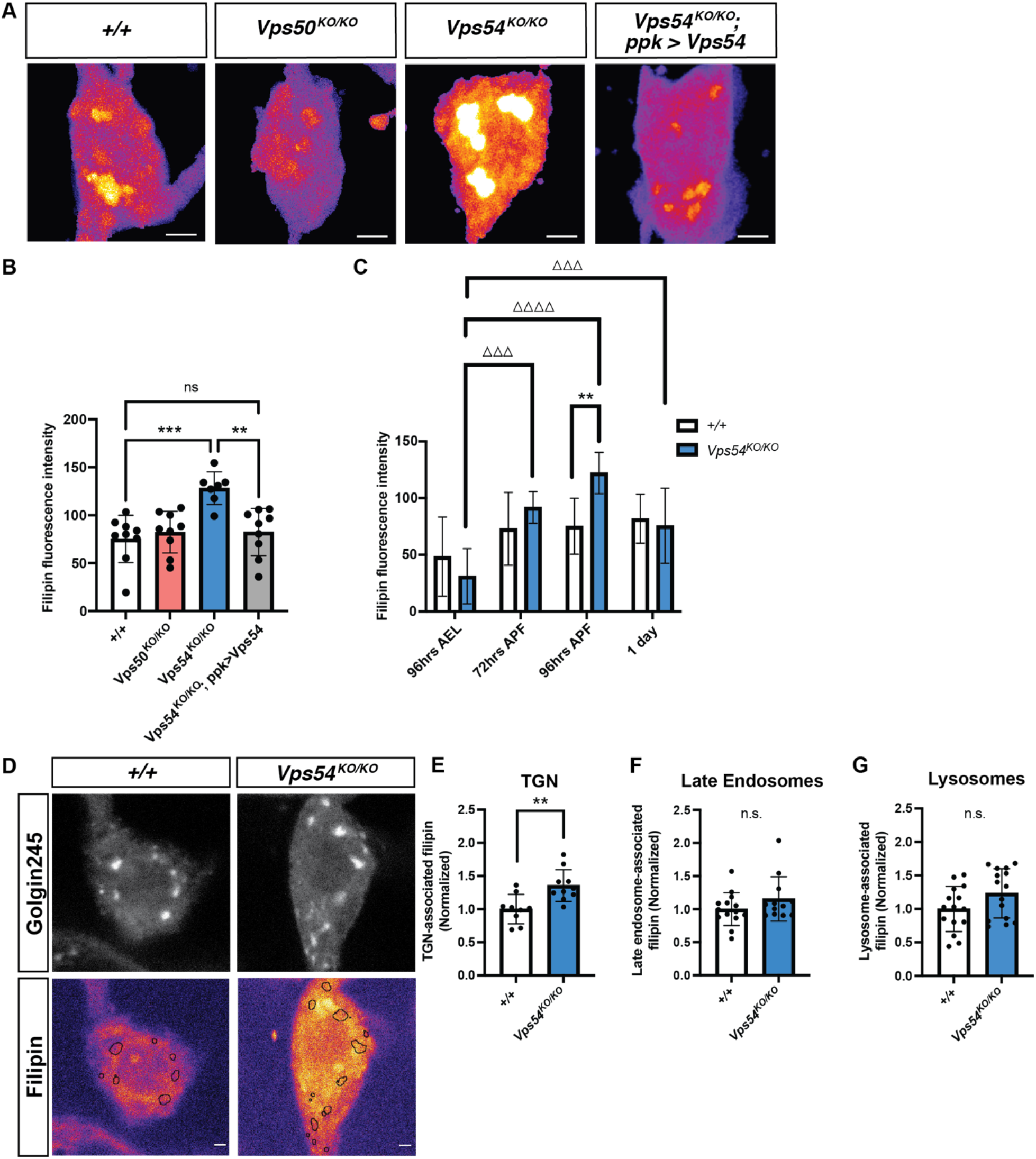
Accumulation of free sterol at the TGN during dendrite regrowth in GARP deficient neurons. A) Maximum intensity projections showing filipin staining in *+/+, Vps50*^*KO/KO*^, *Vps54*^*KO/KO*^, and *Vps54*^*KO/KO*^; *ppk>Vps54* neurons at 96hrs APF. Scale bar = 2.5μm. B) Quantification of filipin fluorescence intensity at 96hrs APF, n = 7-11/genotype. Data normalized to average control value for each experiment to account for inter-experimental differences in filipin intensity. Analyzed by one-way ANOVA with Tukey’s post-test. C) Quantification of filipin fluorescence intensity in *+/+* and *Vps54*^*KO/KO*^ neurons across development. Analyzed by two-way ANOVA with Šidák’s multiple comparison’s correction. ** p< 0.001 for comparison of +/+ to *Vps54*^*KO/KO*^. ΔΔΔ p < 0.001, ΔΔΔΔ p < 0.0001 for comparison of *Vps54*^*KO/KO*^ over time. D). Single plane confocal images of Golgin245 (top) and filipin staining (bottom). Scale bar = 1μm. Quantification of E) TGN-associated, F) late endosomal-associated, and G) lysosomal-associated filipin.

A previous study in mammalian cells determined that sterols accumulate in the late endosomal/lysosomal compartment in Vps52-deficient cells due to missorting of NPC2 (Wei *et al*., 2017). Surprisingly, we did not observe any significant accumulation of sterols in endolysosomes labeled with either Rab7 or Spinster-RFP (Fig. 6, f and g). While we observed a strong filipin signal in the ER as indicated by the marker Sec61β, we did not find any differences in filipin intensity between genotypes in this organelle (Fig. S4, b and c). Of the organelle markers we examined, we only found a significant increase in filipin staining in the Golgin245-positive compartment corresponding to the TGN (Fig. 6, d and e). It thus appears that in GARP-deficient neurons, sterols are capable of exiting the endolysosomal pathway but aberrantly accumulate in the secretory pathway instead.

### Targeting specific lipid regulators at the TGN modulates GARP KO phenotypes

To gain a better understanding of how sterols may be accumulating at the TGN in the *Vps54*^*KO/KO*^, we examined genetic interactions between *Vps54* and various sterol and lipid regulatory proteins. Oxysterol binding protein (Osbp) regulates transport of sterols across several interorganelle contact sites. At contacts between the ER and TGN, Osbp interacts with the ER-localized protein VAP-A to facilitate the transfer of sterol from the ER in exchange for PI4P (Mesmin *et al*., 2017). We therefore made crosses to bring either a null Osbp allele (*Osbp*^*1*^) (Ma, Liu and Huang, 2010) or UAS-Osbp into the *Vps54*^*KO/KO*^ background to decrease or increase Osbp levels, respectively. Removal of one functional Osbp allele (*Vps54*^*KO/KO*^; *Osbp*^*1/+*^) rescued the dendrite morphology defect in *Vps54*^*KO/KO*^ neurons, while Osbp overexpression (*Vps54*^*KO/KO*^; ppk>Osbp) dramatically exacerbated it (Fig. 7, a and b). To evaluate the contribution of Osbp acting at TGN/ER contact sites, we next targeted the PI4-kinase that phosphorylates phosphatidylinositol to generate PI4P, the kinase known as *four wheel drive (fwd*) (Polevoy *et al*., 2009) in *Drosophila*. We reasoned that if Osbp-mediated exchange of sterol for PI4P between the ER and TGN was responsible for the accumulation of sterol at the TGN in *Vps54*^*KO/KO*^ neurons, then *fwd* knockdown should rescue the *Vps54*^*KO/KO*^ dendrite morphology defect. However, expressing a *fwd* RNAi in *Vps54*^*KO/KO*^ neurons exacerbated the *Vps54*^*KO/KO*^ phenotype (Fig. 7, a and b). To further look into the possibility that TGN/ER contacts were responsible for the sterol accumulation in *Vps54*^*KO/KO*^ neurons, we examined interactions between *Vps54* and the single *Drosophila* homolog of VAP-A/B, *Vap33* (Pennetta *et al*., 2002). Neither knockdown nor overexpression of Vap33 had any effect on the *Vps54*^*KO/KO*^ phenotype (Fig. S5). Taken together, these results suggest that TGN/ER contacts are unlikely the source for sterol accumulation at the TGN in *Vps54*^*KO/KO*^ neurons.

**Figure 7.**
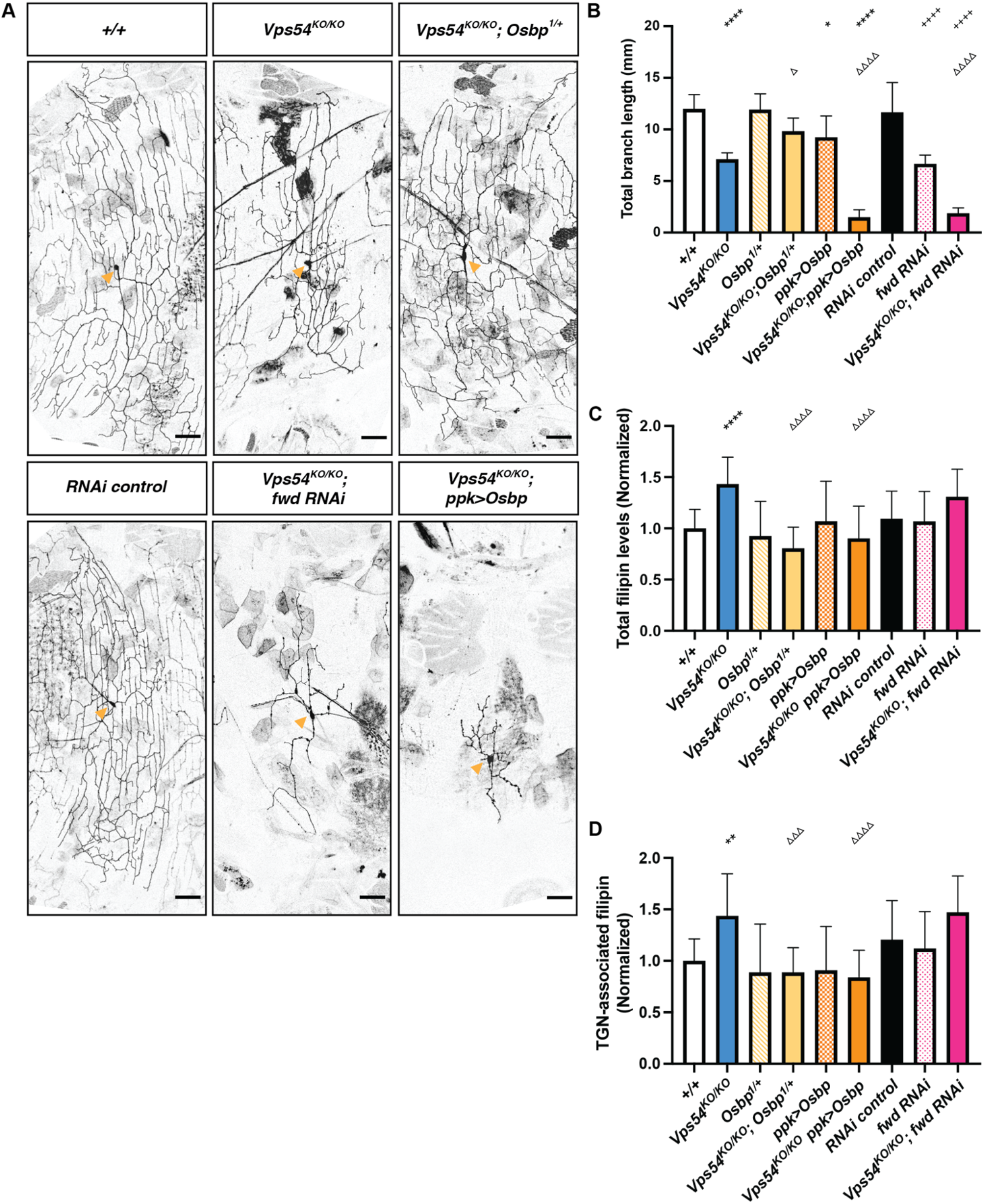
Targeting specific lipid regulators at the TGN modulates GARP KO phenotypes. A) Representative maximum z-projections of class IV da neurons from 7 day old males. Scale bar = 2.5μm Yellow arrows point to the soma. B) Quantification of total dendrite branch length, n > 10 except RNAi control (n = 8), *+/+* and *Vps54*^*KO/KO*^ (both n = 6). Quantification of C) total filipin fluorescence intensity and D) TGN-associated filipin levels at 96hrs APF. For C & D, n > 10 neurons. Analyzed by one-way ANOVA with Tukey’s post-test. * indicates a significant difference from *+/+*, Δ indicates a significant difference from *Vp54*^*KO/KO*^, and + indicates a significant difference from RNAi control. Not all pairwise comparisons shown.

We also examined the effect of targeting *Osbp* and *fwd* on filipin staining in *Vps54*^*KO/KO*^ neurons (Fig 7, c and d). Both total and TGN-associated filipin levels were decreased to control levels when *Osbp* levels were decreased in *Vps54*^*KO/KO*^ neurons (*Vps54*^*KO/KO*^; *Osbp*^*1/+*^), as compared to *Vps54*^*KO/KO*^ alone. Unexpectedly, overexpression of Osbp in the *Vps54*^*KO/KO*^ background also decreased both total and TGN-associated filipin levels, despite exacerbating the dendritic phenotype. When *fwd* was knocked down (*Vps54*^*KO/KO*^; *fwd RNAi*), total and TGN-associated filipin levels remained elevated and were not significantly different from those in *Vps54*^*KO/KO*^ neurons. These data together suggest that Osbp-dependent sterol transfer sites(s) other than the sterol/PI4P exchange cycle between the TGN and ER must contribute to the elevated sterol levels at the TGN.

To further investigate the ability of a single *Osbp*^*1*^ allele to rescue the *Vps54*^*KO/KO*^ phenotypes, we also examined organelle morphology. *Osbp*^*1*^ heterozygosity rescued the number of Golgin245 puncta, but not the number of Rab7+ late endosomes in *Vps54*^*KO/KO*^ neurons (*Vps54*^*KO/KO*^; *Osbp*^*1/+*^) (Fig 8). These results further support the notion that the inability of dendrites to regrow in *Vps54*^*KO/KO*^ neurons is due to perturbations at the TGN, and not to impaired endolysosomal trafficking.

**Figure 8.**
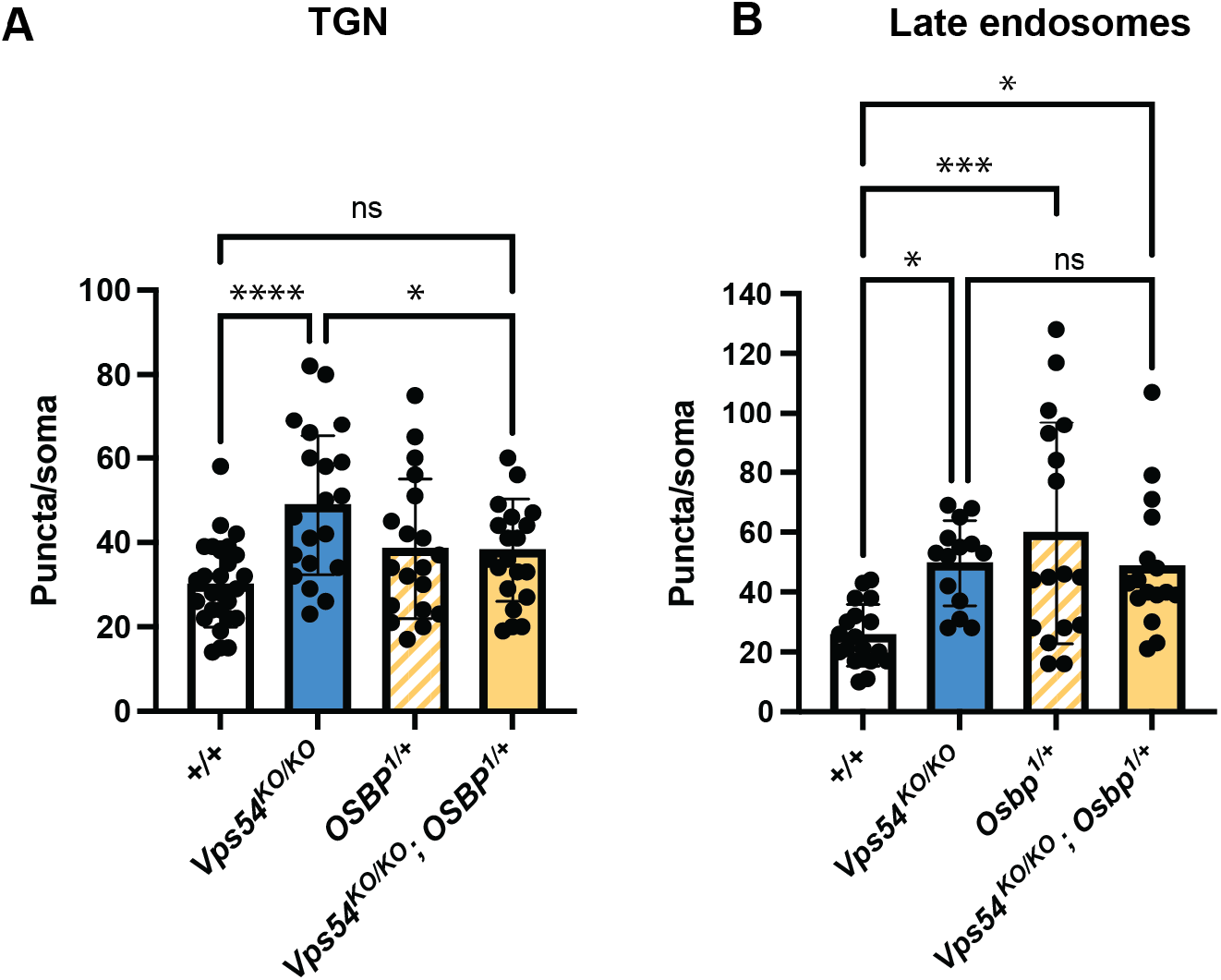
Depletion of Osbp rescues TGN but not late endosomal morphology in GARP KO neurons. Quantification of the number of A) Golgin245 puncta and B) Rab7 puncta per soma in neurons at 96hrs APF. Analyzed by one-way ANOVA with Tukey’s post-test. * p<0.05, ** p<0.01, *** p< 0.001.

## Discussion

### GARP and EARP in neurodevelopment

Despite their links to neurodevelopmental disease, our understanding of how the GARP and EARP complexes function in neurons remains limited. Studies of these complexes have been hampered by the early embryonic lethality of mice lacking components of these complexes (Schmitt-John *et al*., 2005; Sugimoto *et al*., 2012). In our *Drosophila* mutant studies, we show that *Vps50, Vps53* or *Vps54* is dispensable for larval development. Loss of *Vps53* resulted in pupal lethality, while *Vps54* knockouts had a reduced lifespan as adults. Both the GARP and EARP complexes are required for dendrite regrowth in c4da neurons after developmental pruning in pupae. The emergence of this phenotype only at later developmental stages is reminiscent of the secondary microcephaly that emerges postnatally in patients with GARP/EARP complex mutations (Feinstein *et al*., 2014; Gershlick *et al*., 2018; Hady-Cohen *et al*., 2018).

We did not detect gross morphological changes in axon projections, but we cannot rule out finer structural changes, or later degeneration that may occur as a result of GARP/EARP complex deficiency. Given that we observed a reduced lifespan in the *Vps54*^*KO/KO*^ flies, it will be of interest to examine in future studies whether age-dependent changes in neuronal morphology or function may occur in addition to the developmental phenotypes characterized in this study. Studies in the wobbler mouse, bearing a spontaneous point mutation in *Vps54*, reveal degeneration of multiple brain regions and motor neurons in adult mice (Schmitt-John *et al*., 2005; Schmitt-John, 2015). The motor neuron phenotype in mice is distinct from that observed in *Drosophila* mutants harboring the *Vps54* null allele, *scat*^*1*^, which exhibit overgrowth of the larval neuromuscular junction (Patel *et al*., 2020). Taken together with our study, these findings suggest that the effect of GARP/EARP deficiency may be somewhat context or cell-type-dependent. What is clear is that neurons are sensitive to the loss of these protein complexes.

### Appearance of endolysosomal phenotypes only in later developmental stages

At the subcellular level, loss of either the GARP or EARP complexes results in distinct effects on the endolysosomal system. Knockout of *Vps50* specifically affects early endosomes, while knockout of *Vps54* specifically affects late endosomes and lysosomes. Previous studies have shown that the GARP complex is essential for the proper sorting of lysosomal hydrolases. Despite an enlargement of the lysosomal population, we detected no changes in expression or maturation of the hydrolase cathepsin L in head lysates from 1 day old knockout flies. This is consistent with a report on the retromer complex, which functions upstream of the GARP complex in hydrolase sorting. In that study (Ye *et al*., 2020), changes in cathepsin L processing are observed in 30 day old, but not 1 day old, *Vps29* mutant flies. The authors of that study suggest that there must be compensatory mechanisms that facilitate proper lysosomal hydrolase sorting during earlier stages of development. Our results showing that cathepsin L processing is intact in young adult flies, as well as the dispensable role of the GARP/EARP complexes for overall larval development further support this notion.

### GARP in sterol transport in neurons

In *Saccharomyces cerevisiae, Vps53Δ* or *Vps54Δ* cells accumulate sterol intracellularly (Fröhlich *et al*., 2015). As yeast lack the EARP complex, this was assumed to be a function of the GARP complex. In mammalian cells, knockdown of the shared component Vps52 results in mis-sorting of NPC2 (Wei *et al*., 2017), leading to sterol accumulation in lysosomes. This study, however, did not target the complex specific components of the GARP and EARP complexes. We find that sterol accumulates in neurons of *Vps54*^*KO/KO*^ but not *Vps50*^*KO/KO*^ *Drosophila*, suggesting that the EARP complex may not be involved sterol transport. To our surprise, we observed accumulation of sterol in the *Vps54*^*KO/KO*^ neurons at the TGN, not in lysosomes. Osbp likely facilitates sterol transport to the TGN in *Vps54*^*KO/KO*^ neurons, as reducing Osbp levels with a single null allele (*Vps54*^*KO/KO*^; Osbp ^1/+^) rescued sterol levels, TGN morphology, and the dendritic phenotype of GARP deficient neurons. Strikingly, *Osbp*^*1/+*^ heterozygosity did not rescue the observed defects in endolysosomal morphology, indicating that these changes are still permissive to dendrite regrowth. Taken together with the data showing no impairment of cathepsin L maturation in *Vps54*^*KO/KO*^ lysates, our results suggest that perturbed dynamics at the TGN, but not in endolysosomes, contributes in part to the impaired dendrite regrowth.

Our results showing that overexpression of Osbp in *Vps54*^*KO/KO*^ neurons also decreased filipin levels at the TGN while exacerbating the dendritic phenotype suggests that this manipulation may disturb the balance of sterol transport at several interorganelle contact sites. For example, it is possible that the decrease in TGN-associated filipin staining upon Osbp overexpression may be an indirect effect of increased transport of sterol out of the secretory pathway through ER-endolysosome contacts (Dong *et al*., 2016). Additionally, Osbp functions that are independent of sterol transport may be influence dendrite morphology. This is supported by our data showing that overexpression of Osbp in the wildtype background decreased total dendrite length without affecting sterol levels. In this context, Osbp overexpression may alter signaling pathways that act in parallel to regulate dendrite morphology. For example, Osbp creates a scaffold for protein phosphatases, including protein phosphatase 2a (PP2A)(Wang, 2005), which is essential for proper dendrite pruning and cytoskeletal dynamics in c4da neurons (Rui *et al*., 2020; Wolterhoff *et al*., 2020).

Because Osbp regulates sterol transport through multiple interorganelle contact sites, further study is required to identify the precise sites involved in the transport of sterol to the TGN in GARP deficient neurons. Our genetic interaction studies with *Vap33* indicate that interorganelle contact sites other than the ER-TGN contact sites mediated by Osbp may be responsible for the accumulation of sterol at the TGN. One site of interest is the TGN-Rab11^+^ recycling endosome contact site. At these sites, Osbp binds the Rab11 interacting protein RELCH (Sobajima *et al*., 2018). This study demonstrated that knockdown of Osbp, Rab11 or RELCH decreased sterol transport to the TGN. Whether these contact sites exist in *Drosophila* is not yet clear as there is no obvious RELCH homolog. However, Rab11 has been shown to colocalize with TGN markers and to facilitate post-Golgi trafficking during photoreceptor development in flies (Satoh *et al*., 2005). Further, fwd can bind to Rab11, thereby localizing recycling endosomes with Golgi structures during cytokinesis (Polevoy *et al*., 2009), though it is not clear if this interaction permits sterol transfer. These studies suggest an intriguing hypothesis that an increase in TGN-Rab11^+^ recycling endosome contacts in *Vps54*^*KO/KO*^ neurons may lead to sterol overloading, and further, that these interactions may disrupt post-Golgi secretory trafficking necessary for dendrite regrowth.

As sterol auxotrophs, *Drosophila* may utilize multiple pathways to transfer sterol from endolysosomes to the secretory pathway as they are unable to synthesize sterols endogenously. However, beyond the coordinated function of Npc1 and 2, other routes for sterol egress from endolysosomes remain poorly understood. Several recent studies in mammalian cells have focused on identifying mechanisms for sterol metabolism, and sterol transport specifically, in control cells and/or in cells in which Npc1 is either genetically or pharmacologically inhibited (Scott *et al*., 2015; Trinh *et al*., 2020; van den Boomen *et al*., 2020; Lu *et al*., 2021). We suggest that, given their unique reliance on dietary sterol, *Drosophila* is an ideal model in which to study mechanisms of sterol egress from the endolysosomal pathway. It will be important to conduct future studies to identify the mechanisms of sterol uptake and transport to the TGN in the absence of the GARP complex.

## Methods

### Fly stocks

Flies were reared at 25°C in density-controlled vials containing standard cornmeal-molasses food. Transgenic fly stocks used in this study: C4da neurons were visualized using the ppk-Gal4, UAS-CD4-tdTomato or UAS-CD4-tdGFP lines (Han, Jan and Jan, 2011). The following fly lines were purchased from the Bloomington Stock Center: Chromosomal deficiencies deleting regions around the genes of interest: stock# 24372 (Vps50 Df: Df(2R)BSC348/CyO); stock# 7895 (Vps51 Df: Df(2R)Exel7158/CyO); stock# 27381 (Vps52 Df: Df(2L)BSC810/SM6a); stock# 23680 (Vps53 Df: Df(2L)BSC295); and stock# 7813 (Vps54 Df: Df(2L)Exel8022. RNAi lines: stock# 35787 (RNAi control UAS-mCherry in the VALIUM10 vector), stock# 50548 (Vps51 RNAi), stock# 27985 (Vps52 RNAi), stock# 38267 (Vps53 RNAi), stock# 38994 (Vps54 RNAi), stock# 35257 (fwd RNAi), stock# 27312 (Vap33 RNAi). Other mutant and UAS lines: stock# 26693 (UAS-Vap-33-1), stock# 57348 (*Osbp*^*1*^), stock# 57346 (UAS-Osbp), stock# 42716 (UAS-spinster-RFP), stock# 64747 20XUAS-tdTomato-sec61β. The following RNAi lines were purchased from the Vienna Drosophila Resource Center: stock #60200 (KK RNAi control), stock 108290 (Vps50 RNAi).

### Molecular cloning

To generate the UAS-Vps50-3xHA line, Vps50 cDNA was amplified from DGRC clone FI23003. Restriction sites and a C-terminal 3xHA tag were added during amplification (primers listed in Supplemental Methods Table S1). The resulting amplification product was cloned into the Not and Kpn restriction sites in the pACU backbone. To generate the UAS-Vps53-3xHA line, Vps53 cDNA was amplified from DGRC clone clone FI1784. AttB sites were added during amplification. The amplification product was subsequently cloned in the pDONR-221 entry vector using BP-Clonase II (Invitrogen) and subsequently transferred to the pTWH vector (DGRC clone 1100) using LR-Clonase II (Invitrogen). The UAS-Scat line was generated using the expression ready pDNR-Dual-UAS-Scattered-Flag-HA plasmid (DGRC clone FMO06004).

The UAS-Vps50 plasmid was injected into the VK20 attP docking for φC-31-mediated integration (Bateman, Lee and Wu, 2006) site by Genetivision. The UAS-Vps53 and UAS-Scat plasmids were injected into Bloomington stock 24866 (M{vas-int.Dm}ZH-2A, PBac{y[+]-attP-9A}VK00019) by Rainbow Transgenics.

### Generation of knockout flies by CRISPR

Knockout flies were generated by CRISPR/CAS9 homology-dependent repair in which the gene of interest was replaced by an eye-specific dsRed cassette. (See Fig S1 for schematic). Guide RNA sequences were designed using the Fly CRISPR target finder (http://flycrispr.molbio.wisc.edu/tools). Selected sequences were in the 5’UTR and 3’UTR of the gene of interest. Each guide RNA was cloned into the pU6-BbsI-chiRNA. Guide RNA sequences and genotyping primers can be found in Table S1.

The donor template was generated by cloning homology arms (∼1kB upstream of the 5’ guide RNA sequence and ∼1kB downstream of the 3’ guide RNA sequence, see Table S1 for primer sequences) into the pHD-dsRed-attB plasmid. For Vps50 and Vps53, 5’ homology arms were cloned into the NotI site, while 3’ homology arms were cloned into the SpeI site. For Vps54 (scat), the 5’ homology arm was cloned into the AarI site, while 3’ homology arm was cloned into the SapI site.

Guide RNA plasmids and donor plasmids were injected into isogenized *vasa*-cas9 flies (Rainbow Transgenics). DsRed^+^ flies were selected and crossed to balancers. DNA isolated from homozygous DsRed^+^ flies was used for initial genotyping. To generate a control line, the isogenized vasa-cas9 flies were treated in the same manner as DsRed^+^ flies. The resultant line was used as the control (+/+) unless otherwise indicated.

After initial screening, the DsRed cassette was removed by crossing to flies expressing Cre recombinase (specific line). DsRed-offspring from this cross were mated to a second chromosome balancer line. Homozygous progeny were used for genotyping to confirm the absence of the gene of interest (Fig S1).

### RT-PCR

Total RNA was isolated from wandering third instar larvae by TRIzol/chloroform extraction and treated with the TURBO DNA-free reagent (ThermoFisher) to remove genomic DNA. The High-Capacity cDNA Reverse Transcription Kit (Applied Biosystems) was used for cDNA synthesis. Primer sequences for *Vps50, Vps53, Vps54* and three internal controls can be found in Table S1. PCR reactions were performed on equal amounts of cDNA in a with SYBR Green PCR Master Mix (Applied Biosystems). PCR products were run on 1% agarose gels with the GeneRuler 1kb Plus DNA Ladder (ThermoFisher).

### Lifespan analysis

Lifespan analysis was conducted at 25°C. Groups of 10 age-matched flies were collected as they eclosed and transferred to yeasted vials containing standard cornmeal-molasses food. Every 3-4 days, surviving flies were transferred to fresh vials and the number of dead and surviving flies were recorded. Flies were excluded from the study if they escaped, were accidentally crushed or were stuck in food while still alive. Kaplan-Meier curves were generated in Prism (GraphPad) and analyzed by Mantel-Cox log rank test with Bonferroni correction for multiple comparisons.

### Antibodies

The following antibodies were used in this study: anti-rab7 hybridoma supernatant (1:5) and anti-golgin245 (both developed by S. Munroe and obtained from the Developmental Studies Hybridoma Bank). Anti-rab5 (1: 250, Abcam Ab21261). Anti-tdTomato (1:500, Kerafast EST203). Anti-cathepsin L antibody (1:500, R&D Systems MAB22591). The Anti-tubulin antibody (1:2000, Sigma T9026). The anti-scattered (Vps54) antibody was generated by and obtained from R. Sinka (Fári *et al*., 2016) (1:400). Secondary antibodies for immunohistochemistry were anti-mouse, goat, or rabbit labeled by Alexa 488, 555, or 647 (1:1000, ThermoFisher). Secondary antibodies for western blot were anti-mouse HRP (1:1000, Jackson 115-035-146), anti-guinea pig HRP (1:500, Sigma A5545) or anti-mouse IR-Dye 680 LT (1:20000, LI-COR).

### Western blotting

10-20 whole larvae or 30-40 heads from 1 day old flies were homogenized in 50mM Tris-HCl pH 7.4, 150mM NaCl, 1% Triton X-100, 5mM EDTA, 1mM PMSF and 1x complete protease inhibitor (Roche) using a pestle. Samples were then centrifuged for 10 min at 12,000 x g. Samples for gel electrophoresis were prepared 2x Laemmli Buffer (Bio-Rad 1610737) with 5% β-mercaptoethanol. Lysates were heated at 95°C for 10 min, followed by pulse centrifugation. Samples were loaded on 4-12% Bolt Bis-Tris Plus (ThermoFisher) and run in NuPage MES buffer (ThermoFisher). Proteins were then transferred to Immobilon-FL PVDF membrane (Millipore), blocked in 5% milk in TBST (Tris-buffered saline + 0.1% Tween-20). Primary antibodies were diluted in blocking solution and incubated overnight at 4°C. After washing with TBST, membranes were incubated with secondary antibodies for 2hrs at room temperature. Membranes were then again washed before detection. Vps54/scat protein was detected using HRP secondary antibodies with the SuperSignal West Pico ECL chemiluminescent substrate (ThermoFisher 34580) and scanned on a C-DiGit blot scanner. Cathepsin L western blots were detected using LI-COR secondaries and scanned on the LI-COR Odyssey CLx.

### Imaging dendrite morphology

For larval and pupal imaging, staged embryo collections were performed on yeasted grape agar plates at 25°C. Third instar larvae (96hrs AEL) were anesthetized in ether and whole mounted in glycerol. Staged pupae (72 or 96hrs APF) were dissected from the pupae case and mounted on a custom acrylic disc (de Vault *et al*., 2018) without any anesthesia. For adult imaging, flies that eclosed within an 8hr time window were collected as age-matched adults. Flies were aged at 25°C in yeasted vials and were transferred to fresh vials every 3-4 days. Flies were anesthetized with CO_2_ and whole mounted in glycerol. Z-stacks for dendrite morphology were collected with a 0.5μm z-step on a Leica SP5 laser-scanning confocal microscope equipped with a 20x oil immersion objective and the LAS X acquisition software.

### Immunohistochemistry and filipin staining

Larvae, pupae or adults were filleted and fixed in 4% paraformaldehyde for 20 min followed by permeabilization using 0.5% triton X. Fillet preps were blocked in 10% serum and then incubated with primary antibody while rotating overnight at 4C. After primary antibody was washed away, fillets were incubated with secondary antibody for two hours while rotating at room temperature. Fillets were mounted in Diamond ProLong Anti-fade mounting reagent and imaged on Z-stacks for analysis of organelles or filipin staining were collected with a 0.25μm z-step on a Leica SP8 laser-scanning inverted confocal microscope equipped with a 63x oil immersion objective, 3-6x zoom digital zoom, and the LAS X acquisition software. For information on antibody sources and dilutions used, see Supplemental Methods Table S2. Samples to be stained with filipin were first fixed and then stained with 5μg/mL filipin in PBS for 2hrs at room temperature without permeabilization. If filipin-stained samples were also to be stained with antibodies, samples were then permeabilized and stained using the same procedure described above.

### Image analysis

All image analysis was performed in ImageJ Fiji (http://fiji.sc). Morphological analysis of dendrite arbors was performed on maximum projections of z-stacks. Arbors were reconstructed using the Simple Neurite Tracer (Longair et al. 2011). Total dendrite branch length is the summed length of all dendrite branches from a single neuron reconstruction. Sholl Analysis was performed using the built-in Sholl Analysis function.

For organelle analysis, masks were generated from z-stacks of the tdTomato or tdGFP neuronal membrane marker and applied to z-stacks of organelle staining to isolate organelles in neurons from background (neuronal organelle image). Maximum projections of organelle staining were further processed as 8-bit binary images to create ROI around organelles using the Analyze Particles function. ROIs were then transferred to neuronal organelle maximum intensity projection, and used measure puncta number, area, and mean fluorescence intensity. To measure filipin levels in organelles, a mask was created on the organelle marker z-stack and then applied to z-stacks of filipin staining. Filipin intensity levels were then measured on maximum projections of the masked images.

### Statistical analysis

Statistical analyses were performed in GraphPad Prism software. Survival curves were analyzed by Log-Rank Mantel-Cox test with Bonferroni multiple comparisons correction. Comparison of two genotypes was done by t test, comparison of three or more genotypes was done by one-way ANOVA with Tukey’s multiple comparison’s test. Analysis of two genotypes over time was done by two-way ANOVA with Šidák’s multiple comparison’s correction.

## Supporting information

CEO 2021 Supplemental Materials

## Acknowledgements

We thank Maja Petkovic and Kai Li for critical reading of the manuscript and members of the Jan lab for discussion. We thank Rita Sinka for providing the scattered (Vps54) antibody. This work was supported by a NIH National Institutes of Neurological Disorders and Stroke grant (R35NS097227) to Y.N.J. Both L.Y.J. and Y.N.J are investigators of the Howard Hughes Medical Institute.

