## Supplementary material for "Loss of the GARP but not EARP protein complex drives Golgi sterol overload during dendrite remodeling": CEO 2021 Supplemental Materials

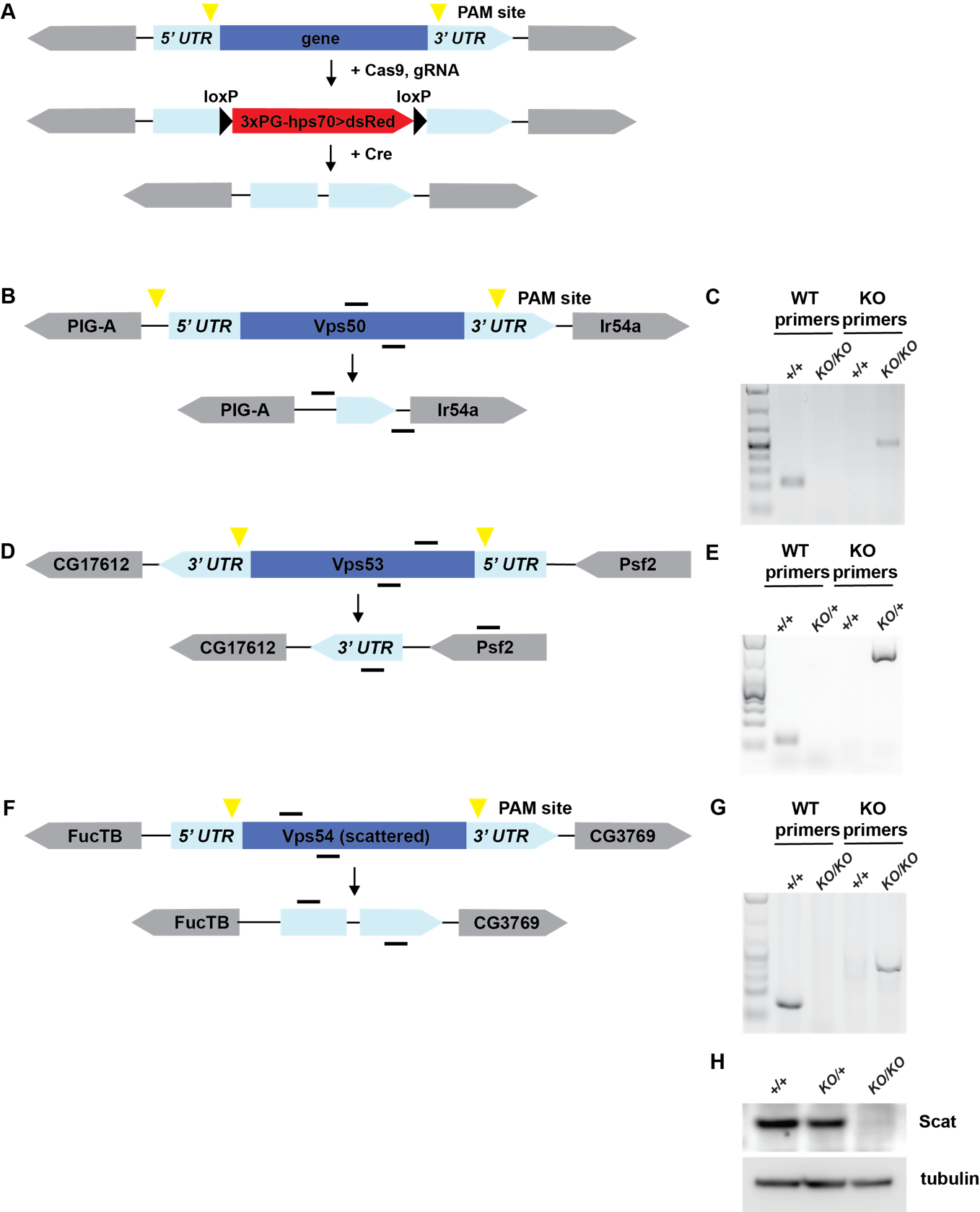

**Figure S1 Generation of GARP and EARP knock-out flies**

A) General schematic of CRISPR knock-out strategy. Guide RNAs recognize sequences around PAM sites (yellow triangles). DsRed cassette flanked by loxP sites was knocked in place of the gene of interest to generate DsRed^+^ knockout (KO) lines. DsRed cassette was removed by crossing to a Cre recombinase line to generate the final knockouts. B, D, and F). Schematic of Vps50, Vps53, and scattered (Vps54) wild type genes and knockouts, respectively. Genes are shown in their relative orientation in the genome. Black lines above indicate hybridization sites for genotyping primers. C, E, G) Agarose gel of genotyping PCR for Vps50, Vps53, and Vps54 knockout lines, respectively. +/+ = w^1118^. For Vps50 and Vps54, DNA was isolated from adult males. Because *Vps53^KO^* is lethal in the pupal stage, DNA was isolated from wandering third instar larvae. H) Western blot of head lysates from control, *Vps54^KO/+^* and *Vps54^KO/KO^* larvae probed with antibodies raised against Vps54/scattered and tubulin (loading control). See Table S1 for cloning and genotyping primer sequences.

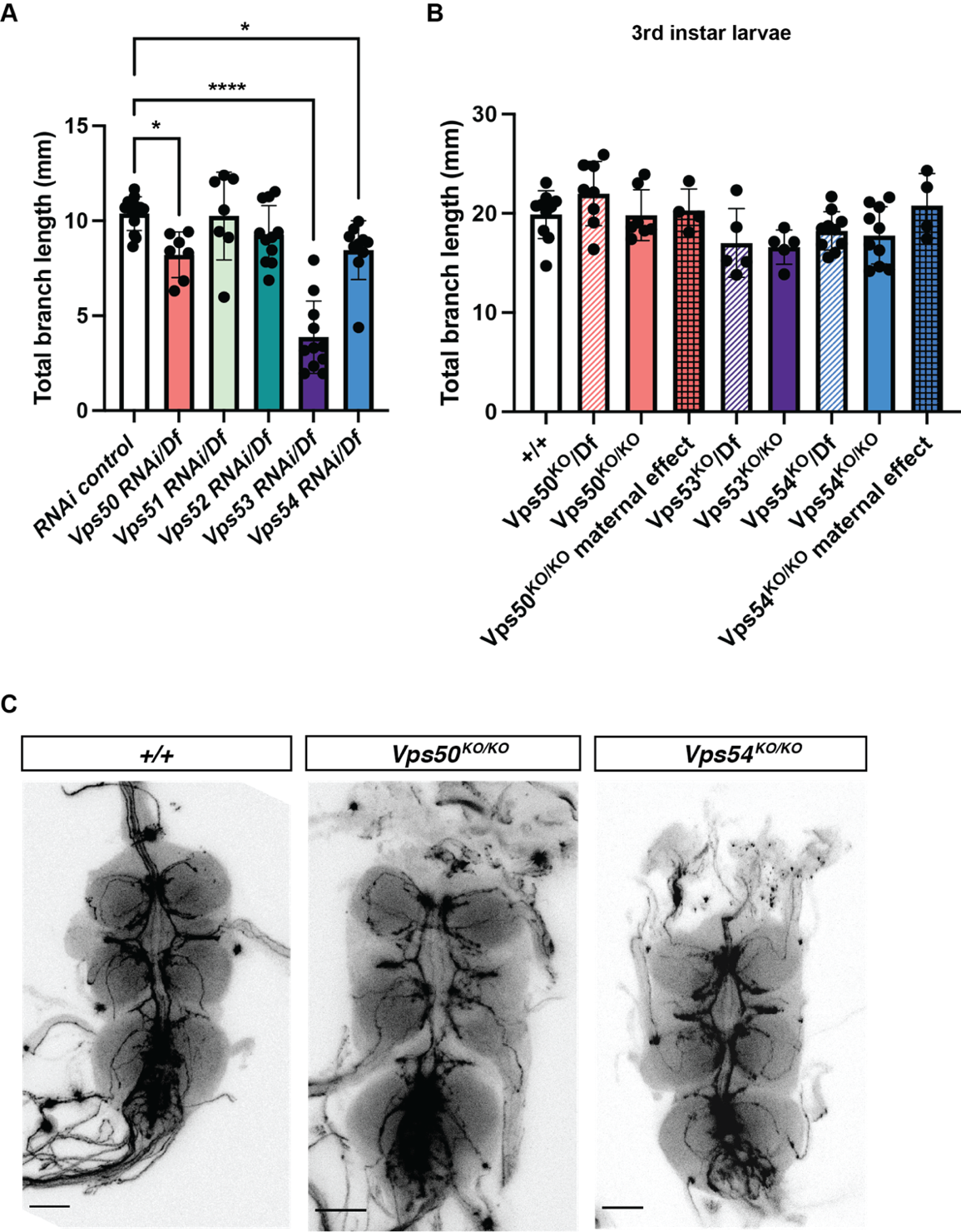

**Figure S2 Additional support for developmental emergence of a dendritic, but not axonal, phenotype** A) Quantification of total dendrite branch length of class IV da neurons expressing RNAi against GARP and EARP complex components in 7 day adult males. RNAi are driven by ppk-Gal4 in a heterozygous chromosomal deficiency background, n = 7-13/genotype. B) Total dendrite branch length of class IV da neurons from 3^rd^ instar larvae (96hrs after egg lay). For maternal effect samples, we examined homozygous KO flies from homozygous KO females crossed to heterozygous KO males. A & B analyzed by one-way ANOVA with Tukey’s post-test. C) Representative images of the ventral nerve cord from 1 day adults. Axons from class IV da neurons are marked with ppk-Gal4, UAS-CD4-tdTomato. Scale bar = 50μm.

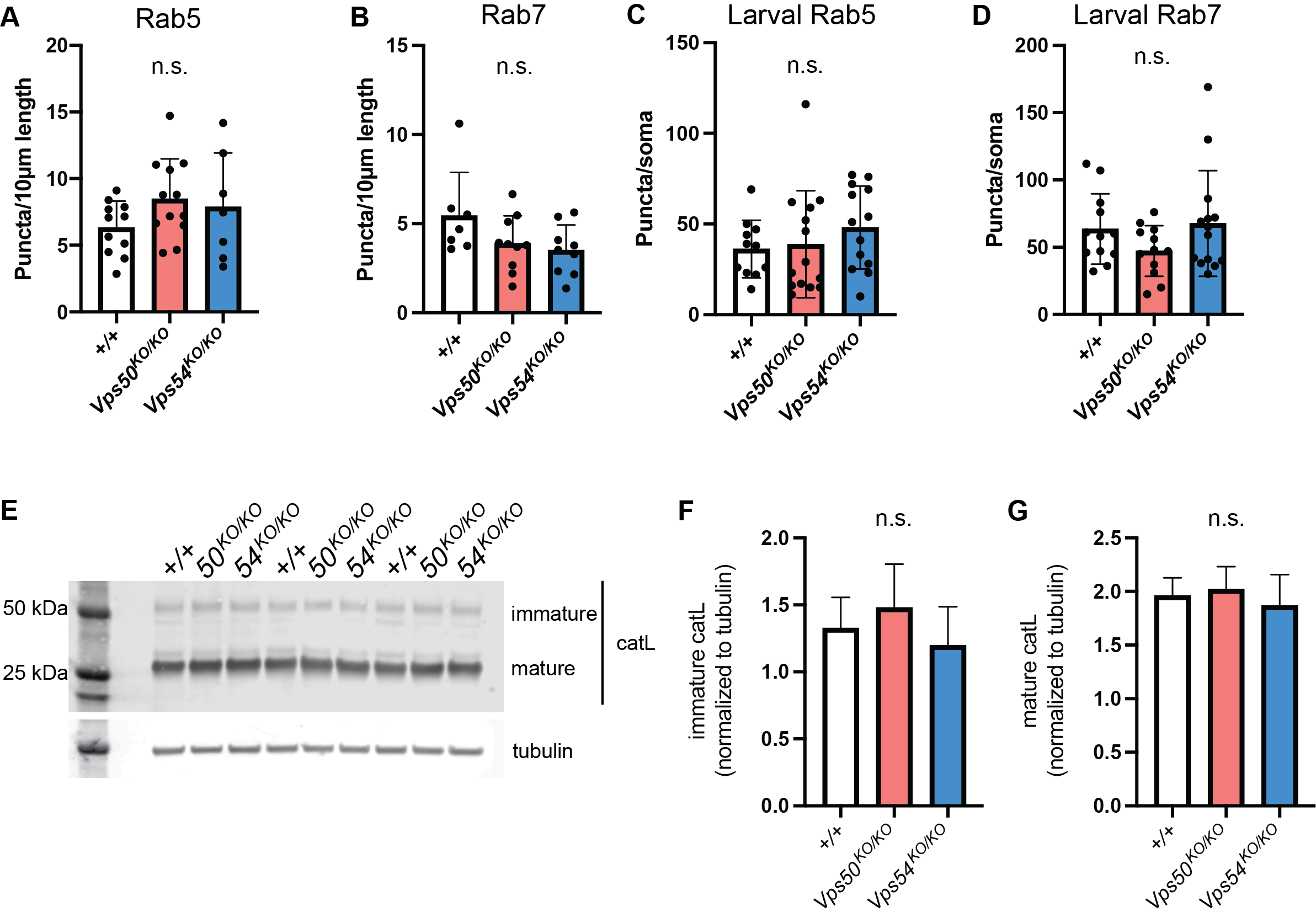

**Figure S3 Endosomal phenotypes in GARP and EARP knockouts**

Quantification of endosomes in proximal dendrites as measured by the number of puncta/10μm of dendrite length in class IV da neurons from 1-day old flies for A) rab5 (n= 7-12) and B) rab7 (n = 7-10). Quantification of puncta in the soma of class IV da neurons from larvae 96hrs after egg lay for C) rab5 (n = 11-15) and D) rab7 (n = 10-14). E) Representative western blot of head lysates from 1 day old *+/+*, *Vps50^KO/KO^* and *Vps54^KO/KO^* flies probed with antibodies against cathepsin L and tubulin. Quantification of the F) immature and G) mature forms of cathepsin L. Analyzed by one-way ANOVA with Tukey’s post-test.

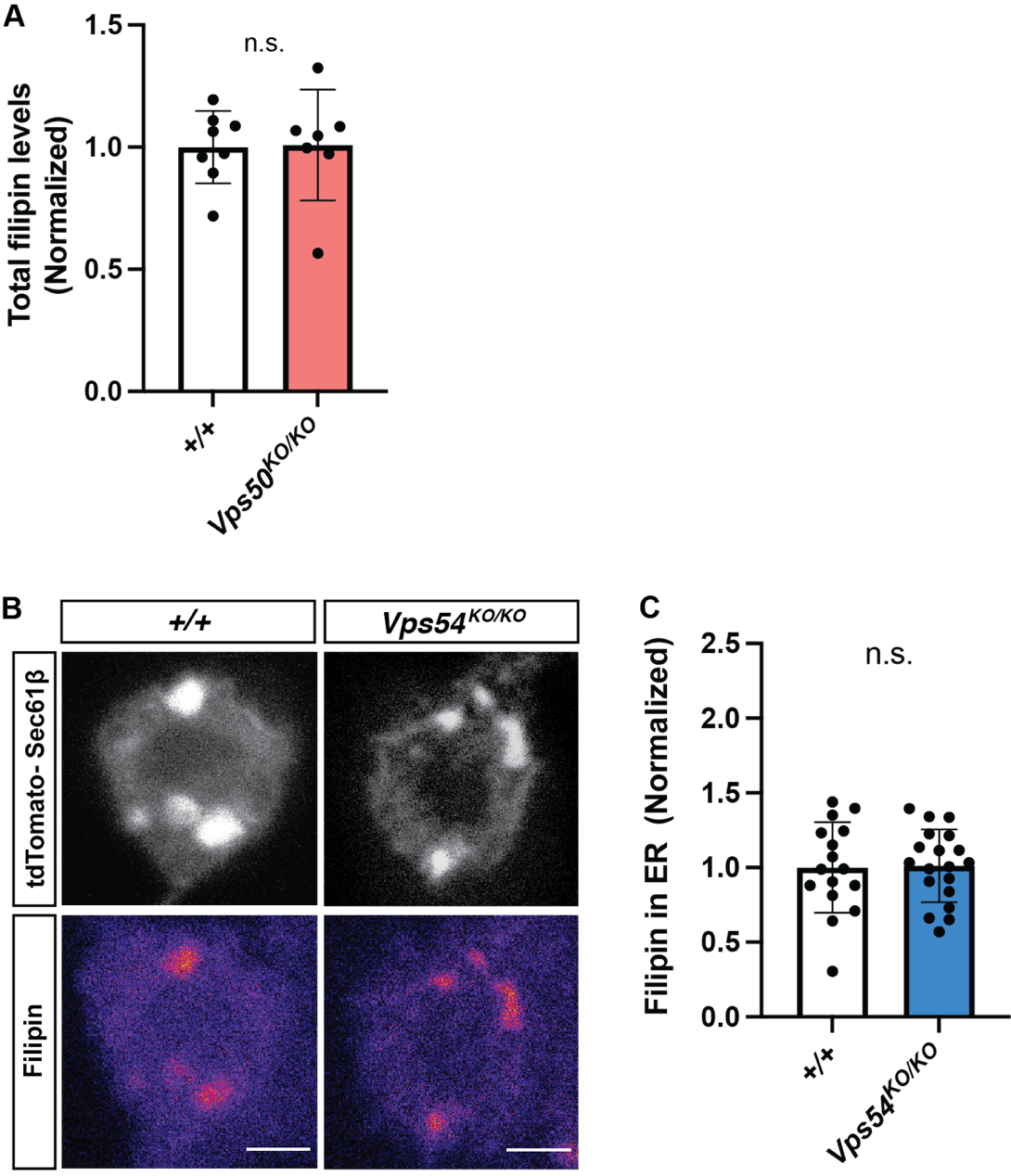

**Figure S4 Sterol accumulation in GARP deficient neurons**

A) Quantification of total filipin levels in neurons from 1 day-old +/+ or *Vps50^KO/KO^* neurons. B) Single plane confocal images of tdTomato-Sec61β (top) and filipin staining (bottom). Scale bar = 2.5μm C) Quantification of ER-associated filipin in +/+ or *Vps54^KO/KO^* neurons. Both A and C analyzed by t test, n.s.

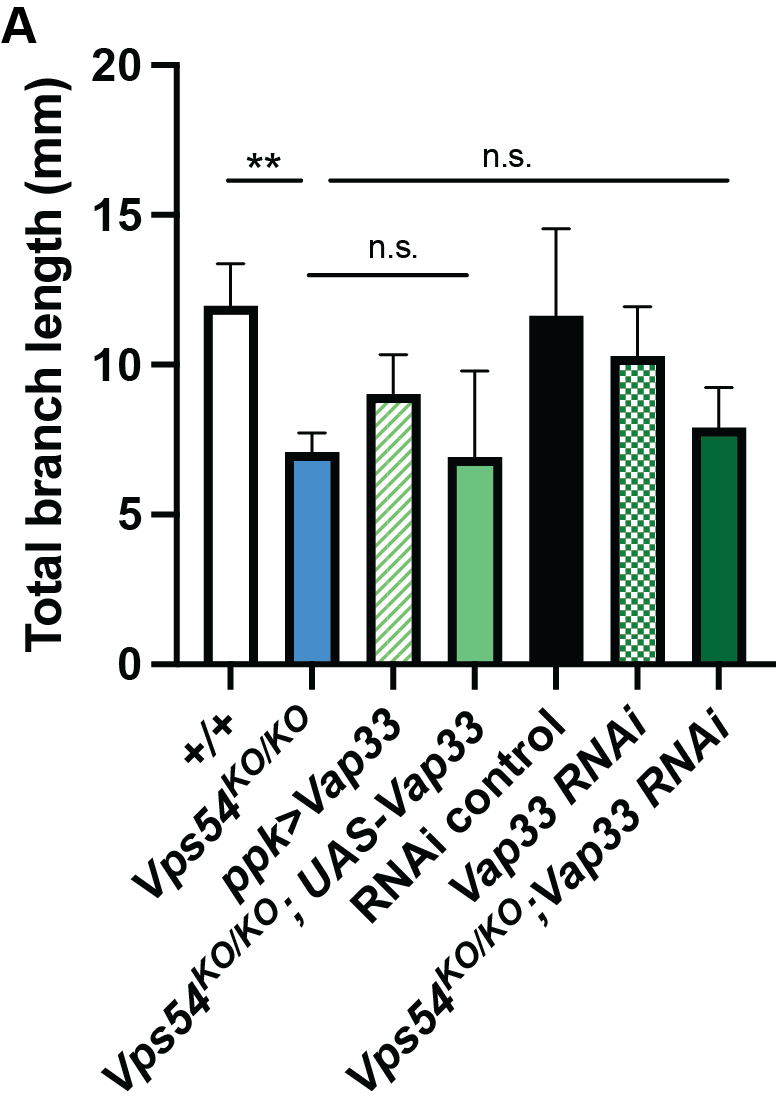

**Figure S5 Changing Vap33 expression does not affect dendrites in *Vps54^KO/KO^* neurons** Quantification of total dendrite branch length of class IV da neurons in 7-day old adults. N = 8-10 for all genotypes except *+/+* and *Vps54^KO/KO^* (n = 6 for both). Analyzed by one-way ANOVA with Tukey’s post-test. ** p <0.01. *Vps54^KO/KO^* is not statistically different from either *Vps54^KO/KO^; ppk>Vap33*, or *Vps54^KO/KO^; Vap33 RNAi*.

**Table S1. Primers used for generating and genotyping knockout flies by CRISPR**

| *UAS plasmids* |  |
| --- | --- |
| Vps50 NotI 5’ | AAAAAGCGGCCGCATGCAGAATTCCAAGGCCAAAATGGA |
| Vps50 3x HA Kpn 3’ | AATTAGGTACCTCAGGTGGACCGGTGTCCGC |
| Vps53 5’ attB | GGGGACAAGTTTGTACAAAAAAGCAGGCTTACACCATGAGCGAGTCTGCG |
| Vps53 3’ attB | GGGGACCACTTTGTACAAGAAAGCTGGGTTGGGGAAGCGCTTTTTCAGTAGG |
| *CRISPR* |  |
| *Guide RNA sequences* | |
| Vps50 5’ guide RNA | CTTCGAGAAAATGCTAATCGGTACT |
| Vps50 3’ guide RNA | CTTCGGTACTTAAGTCATGGTGAA |
| Vps53 5’ guide RNA | CTTCGTAAATTGCTGTCGTGCTGTA |
| Vps53 3’ guide RNA | CTTCGATGTAGCCGTATATTAACT |
| Vps54 5’ guide RNA | CTTCGGCTGTTCGTTCTCTAGCCT |
| Vps54 3’ guide RNA | CTTCGCAATGAAGACGAACTAGGCT |
| *Primers for cloning homology arms* | |
| Vps50 5’ arm F | ATTCGCGGCCGCTGGAATGATGCCAGCTAGCAAAT |
| Vps50 5’ arm R | ATTCGCGGCCGCTTTAAGCACTTTTATACATATTCGTGGC |
| Vps50 3’arm F | AAAAACTAGTCCAAACCCTCCTTAAGTGCAA |
| Vps50 3’ arm R | AAAAACTAGTCGTTCGTCTGTCCACGTAGAG |
| Vps53 5’ arm F | AAAAGCGGCCGCAAAGTTATTGTTATTCTCTTCGTGG |
| Vps53 5’ arm R | AAAAGCGGCCGCTGTCATCACTGGCCGCTC |
| Vps53 3’arm F | AAAAACTAGTGACCTAATAAATTAAGCAACTAATCAT |
| Vps53 3’ arm R | TATAACTAGTGGAAAAGGATGTCTTTCATAGGTGAGT |
| Vps54 5’ arm F | ATTGCACCTGCCATGTCGCAGGGGAAAGTCTCCCGTCTTT |
| Vps54 5’ arm R | GTGTCACCTGCTAATCTACGCCATAACACATGCTCTGCG |
| Vps54 3’arm F | TTACGCTCTTCGTATGGCTCTACTGAGCATTCCGAA |
| Vps54 3’ arm R | CGTAGCTCTTCTGACTATGCGCACTCCAAATCCGTCCA |
| *Primers for genotyping* | |
| Vps50 WT F | TCTCCGACAACTGCTTTGCT |
| Vps50 WT R | CTCGTCCACAATGCCGGATA |
| Vps50 KO F | GGCTGCAGACGTTTTTGTGT |
| Vps50 KO R | CGGGAACTGCTCCAAGTTGA |
| Vps53 WT F | GAGTACGCCTCCAAAGTGCT |
| Vps53 WT R | GCATATCGCGCGTGAGTAAC |
| Vps53 KO F | CGGGTCCGAACAGTAACTCTC |
| Vps53 KO R | TAGGCGGATGGATTGCGAATA |
| Vps54 WT F | ACTTGAGTTCTGTGCCCGAG |
| Vps54 WT R | AGTGACTCAGCTGTTCCTGC |
| Vps54 KO F | GCCTAGAGAACGAACAGCCA |
| Vps54 KO R | ACCGCCATTGATTTGTACGC |
